# Spatial heterogeneity of an ecologically relevant environment accelerates diversification and adaptation

**DOI:** 10.1101/838110

**Authors:** Stineke van Houte, Dan Padfield, Pedro Gomez, Adela M. Lujan, Michael A. Brockhurst, Steve Paterson, Angus Buckling

## Abstract

Spatial heterogeneity is a key driver for the evolution of resource specialists and has been shown to both promote and constrain the rate of adaptation. However, direct empirical support for these evolutionary consequences of spatial heterogeneity comes from simplified laboratory environments. Here we address how spatial structure, through its effect on resource heterogeneity, alters diversification and adaptive evolution of the soil bacterium *Pseudomonas fluorescens* in an ecologically relevant context: soil-based compost. Our data show that environmental heterogeneity can both promote phenotypic diversification and accelerate adaptation. These results suggest that environmental disturbance, which can decrease spatial heterogeneity, may limit diversification and adaptation of microbial populations.

## Introduction

While there is strong theoretical and empirical evidence that spatial variation in resources can promote the evolution and maintenance of diverse resource specialists^1-9^, the impact of spatial heterogeneity on adaptive evolution is more ambiguous. Theoretically, the structured populations associated with spatial heterogeneity can constrain adaptive evolution by both slowing the spread of beneficial mutations^10^ and increasing the role of genetic drift by reducing effective population sizes^11,12^. However, population structure may promote adaptive evolution by allowing greater exploration of adaptive landscapes^13,14^ and spatial heterogeneity can also increase the chance that beneficial mutations will encounter an environment that maximises their fitness effect^15,16^. *In vitro* experimental studies involving bacteria or viruses evolving in nutrient media provide support for both views^3,12,17,18^. This variation in empirical results demonstrates the nuanced effect of *in vitro* experimental conditions, making it crucial to know how spatial heterogeneity impacts adaptive evolution (and indeed diversification) under ecologically relevant conditions. Here, we conduct such an experiment in potting compost^19^. We evolved the soil bacterium *Pseudomonas fluorescens* in compost with its spatial structure intact (representing a heterogeneous environment), or in compost that was mixed with water (representing a homogeneous environment), and measured fitness, phenotypic diversity based on substrate use and population genomic changes.

## Materials and methods

### Strains

*Pseudomonas fluorescens* strain SBW25^20^ was used throughout the study. To generate a genetically marked SBW25 strain expressing β-galactosidase (LacZ), Tn7-mediated transposition was carried out to insert a *lacZ* gene into the *P. fluorescens att*Tn7 genomic location^21^.

### Growth conditions of the evolution experiment in compost

*Pseudomonas fluorescens* SBW25 was grown overnight at 28°C in King’s media B (KB) in an orbital shaker (180 rpm) and then centrifuged for 10 min at 3500 rpm to produce a bacterial pellet, which was resuspended in M9 salts buffer to a final concentration of 10^8^ colony forming units (CFUs)/ml. Following our previous method^22^, six round petri dishes each containing 25 g of twice-autoclaved compost (John Innes no. 2) were inoculated with 5 ml of the *P. fluorescens* suspension (10^8^ CFUs/ml) to give rise to the heterogeneous environment treatment. These compost microcosms were then placed in an environmental chamber at 26°C and 80% relative humidity. For the homogeneous environment treatment, we used six 30-ml glass vials containing 3 g of compost mixed with 9 ml sterile water (compost-wash microcosm), and inoculated each vial with 5 ml of the *P. fluorescens* suspension (10^8^ CFUs/ml). Compost-wash populations were propagated at 28°C in an orbital shaker at 180 rpm. One third of each culture was serially transferred to fresh remaining two thirds of compost and compost–wash approximately every six days during 48 days.

### Sample collection

At 48 days after the start of the experiment, compost samples (2 g) were collected using a sterile spatula and mixed with 10 ml sterile M9 salts buffer and glass beads, and then vortexed for 1 minute. The resultant sample washes from both treatments were diluted in M9 salts buffer, plated onto KB agar and incubated for 2 days at 28°C to determine CFUs per gram of compost. Differences in densities between treatments were tested using a linear model of log10 CFUs g^−1^ compost as the response and adaptation environment (homogeneous *vs.* heterogeneous) as the predictor. From each replicate experiment a subpopulation of 10 bacterial clones were isolated and stored at −80°C in 20% glycerol for further analysis.

### Phenotypic assays

To measure phenotypic diversity (and partition this into different sources of variation) under either homogeneous or heterogeneous conditions, we performed catabolic profiling using Biolog GN2 microplates (Biolog, Hayward CA). To this end, we used the 10 individual colonies isolated from each of the six replicate experiments of each treatment. Each of the bacterial clones was grown individually overnight in KB broth (28°C at 180 rpm). Bacteria were then diluted 1000-fold in M9 salts buffer and incubated for 2h at 28°C to starve the cells. For every clone, each well of a microplate was filled up with 150 μL of culture suspension containing the starved bacteria and incubated at 28°C for 24h, after which optical density was measured at 660 nm as a proxy for bacterial growth using a plate reader (Bio-Tek Ltd). After filtering the number of substrates where no growth was observed (OD_660_ < 0.1), the catabolic profiles of each clone on 91 different substrates were used in downstream analyses.

The analysis of resource use splits the phenotypic variation, V_P_, within a population into genetic variation, V_G_, environmental variation, V_E_, and genotype-by-environment variation, V_GE_. Differences in V_P_, V_G_ and V_GE_ between evolution environments would indicate that changes in spatial heterogeneity result in differences in resource-use diversity. V_P_ was calculated as the average (by taking the mean) Euclidean distance across all pairs of clones^23^, V_G_ as the average variance of clone performance on each substrate^24^, and V_E_ as the average variance of individual clone performance across all substrates. The decomposition of genotype-by-environment variation into responsiveness and inconsistency gives relative measures of the diversity of resource exploitation strategies and niche differentiation (functional diversity)^24,25^. Responsiveness, *R*, indicates differences in environmental variances of genotypes within a community. A high responsiveness value would mean genotypes utilise the same substrates, but they differ in their environmental variance, such that some genotypes perform similarly across all substrates, whereas some have asymmetric performance. In this way, responsiveness measures the diversity of resource exploitation strategies (i.e. specialists *vs.* generalists):

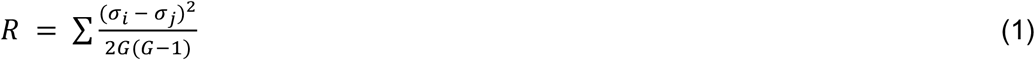

where *G* is the number of genotypes tested within a population and *σ*_*i*_ and *σ*_*j*_ are the standard deviations of environmental responses of each genotype tested. The inconsistency component, *I*, indicates non-correlations between genotypes over environments:

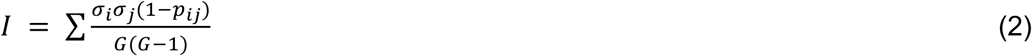

where *ρ*_*ij*_ is the correlation of performance across substrates between each pair of genotypes. High inconsistency suggests that different genotypes within a community have adapted to utilise difference substrates (i.e. different clones perform best on different substrates). In this way, inconsistency is a measure of niche differentiation and the evolution of diversity within populations. In instances of high inconsistency and high responsiveness, genotypes perform best on different substrates, but some of these perform well across many substrates, whereas others perform well on very few. For each variance component, differences between heterogeneous and homogeneous treatments were analysed using linear models, with evolved environment (homogeneous *vs*. heterogeneous) as a predictor compared to a model without any predictor variables.

### Competition assays

Competition assays were performed to evaluate if bacteria evolved in either heterogeneous or homogeneous conditions are better adapted to their own environment. For each microcosm, a mix was generated in which the 10 clones used previously (see above) were pooled together in equal amounts. This mixture was then competed 50:50 with an ancestral *lacZ*-marked *P. fluorescens* clone to allow us to distinguish the mix of evolved clones from the ancestral clone. Competitions were performed in either compost microcosms (heterogeneous competition environment) or in a shaken compost-water mixture (homogeneous competition environment) for 7 days, using a starting inoculum of 10^8^ CFUs total (i.e. 5 × 10^7^ CFUs each of ancestral clone and evolved clone mix). Samples taken at 0 (T0) and 7 (T7) days were diluted in M9 salts buffer and plated on KB agar plates containing 50 μg/ml X-gal to allow blue/white screening. For each microcosm, the numbers of white and blue colonies were used to calculate the relative fitness of each strain: (relative fitness = [(fraction strain A at T7) * (1 – (fraction strain A at T0))] / [(fraction strain A at T0) * (1 – (fraction strain A at T7)])^26^. To look for patterns of local adaptation, we looked at changes in relative fitness with competition (homogeneous *vs.* heterogeneous) and evolution environment (homogeneous *vs.* heterogeneous) added as potentially interacting factors. A linear mixed effects model was used, with population included as random effect to account for the pairing of replicates across treatments. Model selection was done using likelihood ratio tests, and targeted pairwise comparisons were carried out using the *R* package ‘*emmeans*’^27^, where we looked for differences between evolved heterogeneous populations in homogeneous and heterogeneous conditions, evolved homogeneous populations in homogeneous and heterogeneous conditions, and evolved homogeneous populations in homogeneous conditions *vs*. evolved heterogeneous populations in heterogeneous conditions.

### Sequencing

To measure genotypic diversity in clones from each of the treatments, we performed whole genome sequencing (WGS) on pools of the 10 bacterial clones that were isolated from each replicate (pool-seq). In parallel, WGS was carried out on (1) a single clone from each replicate and (2) all 10 individual clones from a single replicate of each treatment. This allowed us to estimate the degree of linkage between mutations for estimating diversity. Each of the 10 bacterial clones were grown individually overnight in KB broth (28°C at 180 rpm). Next day, the cultures were diluted in M9 salts buffer to ensure they had equal densities as measured by OD_600_. Pools of each of the 10 clones were made by mixing equal volumes of each bacterial clone. Total DNA extraction (1.2 ml per sample; 12 pooled-clone samples and 32 single-clone samples) was performed using the Qiagen Blood and Tissue kit following the manufacturer’s instructions. An Illumina HiSeq 2000 sequencer was used to generate 100 bp paired reads from a 500 bp insert library. Reads were trimmed for the presence of Illumina adapter sequences using *Cutadapt* (v1.2.1). The reads were further trimmed using *Sickle* (v1.2) with a minimum window quality score of 20. Reads shorter than 10 bp after trimming were removed. Trimmed reads were mapped to the *P. fluorescens* SBW25 reference with *bwa-mem* (v0.7.12-r1039). For the clonal level sequencing, variants were identified using *GATK Haplotyper* (v3.7) and structural variants were detected using Delly2 (v0.7.7) with a subsequent cut-off of >= 0.95 as a proportion to identify structural variants in haploid genomes. For the pool-seq, sites prone to sequencing or mapping errors were first identified on the clonal ancestor strain using *samtools mpileup* with parameters *-Q0* and *-q0* (i.e. relaxed mapping and base qualities) and then filtered from all subsequent analyses. SNPs were then detected in the pooled populations using *samtools mpileup* with parameters *-Q20* and *-q20* (i.e. relatively strict mapping and base qualities). Indels were identified in pooled data using *scalpel* v0.5.3 (originally designed to detect indels in tumour versus somatic samples^28^) by comparison of evolved with ancestral samples.

### Sequence data analysis

First, we evaluated the ability of our pooled sequencing to correctly identify the number of genetic changes observed in the clonal sequencing (genetic changes with a proportion of >= 0.95). To do this we created a pseudo-pool sequencing file that was based on clonal sequencing where each of the 10 clones from a pool-seq sample had been sequenced individually, such that 10% reads from each file were added into a separate fasta file. These pseudo-pool data were analysed using the same pool-seq pipeline to determine the number of mutations, which should theoretically be equal to the clonal sequencing data (when the cut-off for proportion is >= 0.1). However, whereas 12 genetic changes (8 SNPs and 4 indels) were identified across all the clonal sequencing, for the 2 replicates for which we had sequenced every clone, at least 40 SNPs were identified. With a proportion cut-off of 0.1, we identified SNPs identified in the clonal sequencing, but always identified many more false negatives. It is unclear whether the clonal sequencing underestimates the number of genetic changes, or whether the pool-seq pipeline overestimates such changes. As a result, we took the conservative approach of filtering identified SNPs and indels in the pool-seq data from all the SNPs and indels identified in the clonal sequencing.

We evaluated genetic differences between treatments by calculating (1) the genetic distance of each population from the reference genome, (2) the number of SNPs / indels in each population and (3) alpha diversity, calculated using a modified version of the Hardy-Weinberg equilibrium, such that 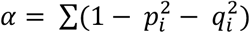, where *i* is the position of each SNP / indel, *p* is the proportion of the SNP / indel and *q* is 1 – *p*. Differences between these metrics were analysed using 2-sample Kruskal-Wallis tests as the data did not conform to the assumption of normality. To test for genetic differences between populations, we performed non-metric multidimensional scaling on the Euclidean distance matrix of SNPs / indels and their proportions in each population using the function ‘*metaMDS*’ in the R package ‘*vegan’*^29^. Permutational ANOVA tests were run using the ‘*adonis*’ function, with Euclidean distance as the response term and evolution environment (homogeneous or heterogeneous) as the predictor variables with 9999 iterations. Changes in beta-diversity were examined using the ‘*betadisper*’ function with the same response and predictor variables in the PERMANOVA.

## Results

We evolved six replicate populations of the soil bacterium *Pseudomonas fluorescens* SBW25 in sterile potting compost (spatially heterogeneous) and a sterile compost-water mix (spatially homogeneous) for 48 days. In this period, populations achieved approximately 3-fold greater densities in the compost-water mix (Fig. 1; likelihood ratio test between models with and without evolution environment as a predictor: *F*_1,10_=14.77, *P* = 0.003).

**Figure 1.**
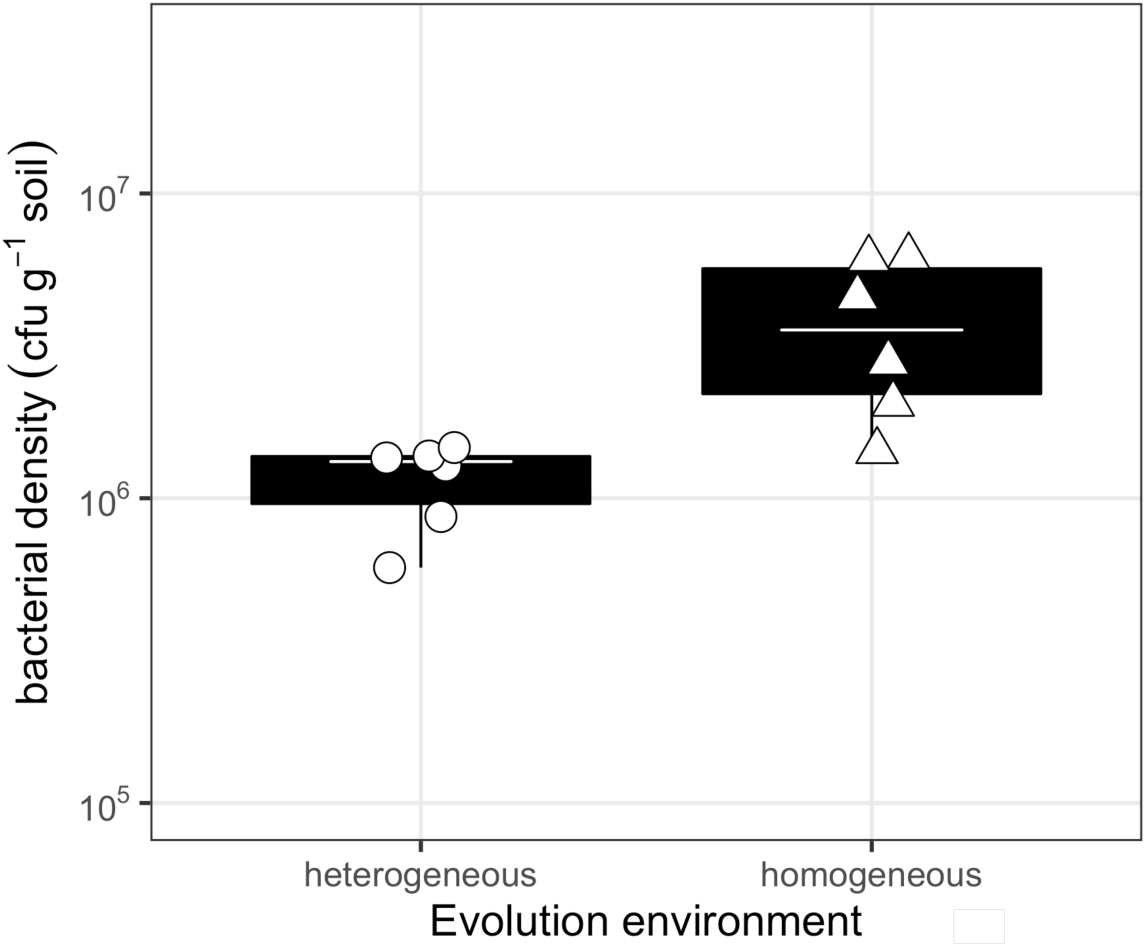
Bacterial densities of populations evolved under either heterogeneous or homogeneous conditions. Points represent densities of each population (CFUs per gram of soil). Tops and bottoms of the bars represent the 75th and 25th percentiles of the data, the white lines are the medians, and the whiskers extend from their respective hinge to the smallest or largest value no further than 1.5 * interquartile range. Points outside this range are outliers.

### Phenotypic data

To test the prediction that spatially heterogeneous environments support the evolution of greater diversification, we isolated 10 individual clones from each replicate population and measured their performance across 96 different substrates (Fig. 2a,b). We then calculated phenotypic variation and partitioned this into V_G_, V_E_ and V_GE_ (Fig. 2c-f). There was no significant impact of environmental heterogeneity on phenotypic variation (likelihood ratio test between models with and without evolution environment as a predictor: *F*_1,10_=2.34, *P* = 0.16) or genotypic variation (likelihood ratio test between models with and without evolution environment as a predictor: *F*_1,10_=2.38, *P* = 0.15; Fig. 2c), but heterogeneous populations did have higher environmental variation (likelihood ratio test between models with and without evolution environment as a predictor: *F*_1,10_=9.131, *P* = 0.012; Fig. 2d). We further decomposed genotype-by-environment variation into responsiveness (measures the diversity of resource exploitation strategies) and inconsistency (a measure of niche differentiation and the evolution of diversity). Responsiveness was not significantly impacted by environmental heterogeneity (likelihood ratio test between models with and without evolution environment as a predictor: *F*_1,10_=4.808, *P* = 0.053; Fig. 2e). However, consistent with a role for spatial heterogeneity in diversification, heterogeneous environments had higher inconsistency (likelihood ratio test between models with and without evolution environment as a predictor: *F*_1,10_=10.026, *P* = 0.010; Fig. 2f) compared to the homogeneous environments. This suggests that heterogeneous environments resulted in higher diversity in resource use than the homogeneous populations.

**Figure 2.**
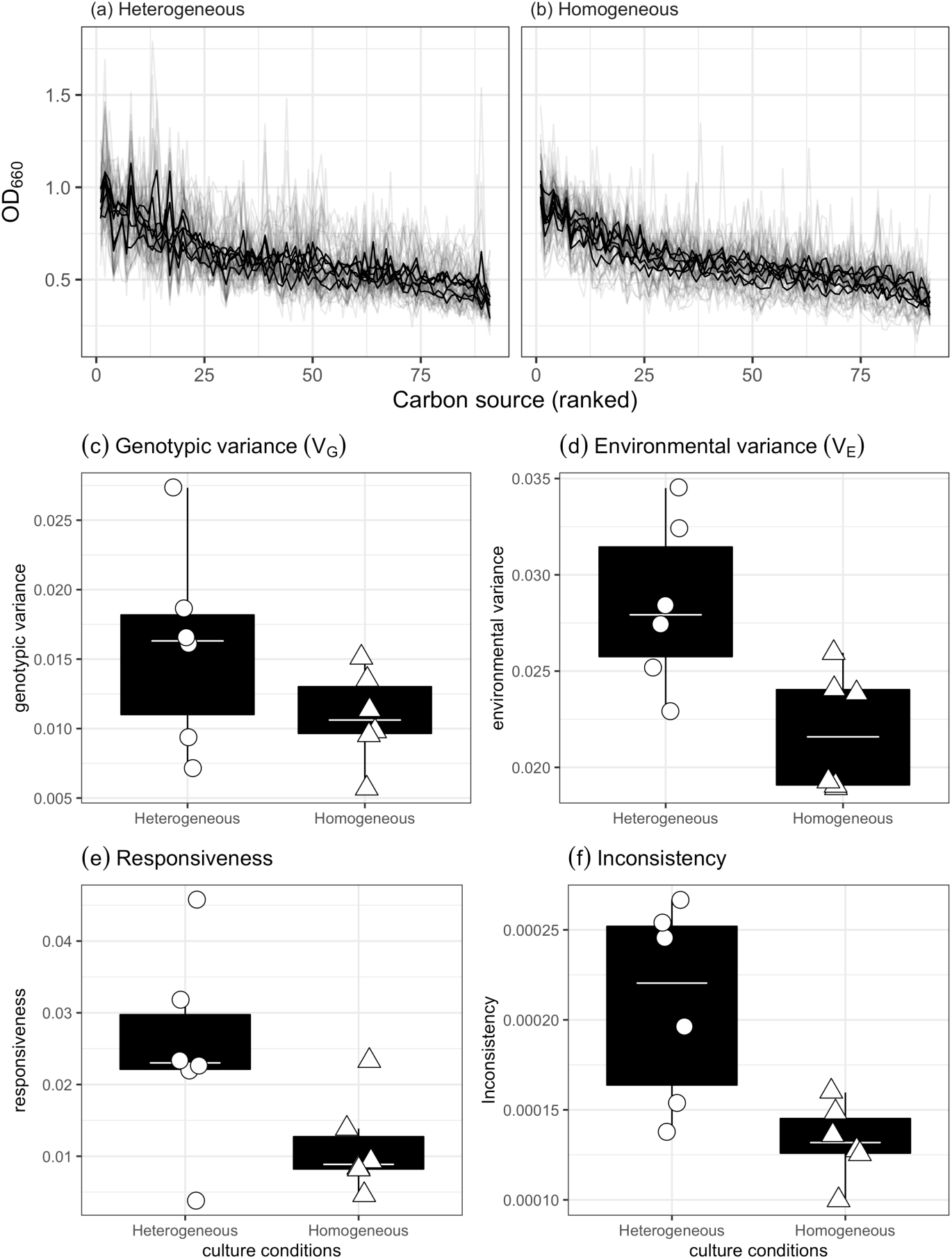
Catabolic profiles of populations evolved under either heterogeneous or homogeneous conditions. Performance of each clone evolved under (a) heterogeneous or (b) homogeneous conditions on a variety of substrates and phenotypic variance was partitioned into (c) genotypic variance (d) environmental variance and genotype × environmental components: (e) responsiveness and (f) inconsistency. Bacteria evolved in a heterogeneous environment had higher environmental variance and higher inconsistency, indicating they had evolved to specialize on different resources. In (a,b), black lines represent the mean OD_660_ of each population and grey lines represent the performance of individual clones. In (c-f) tops and bottoms of the bars represent the 75th and 25th percentiles of the data, the white lines are the medians, and the whiskers extend from their respective hinge to the smallest or largest value no further than 1.5 * interquartile range. Points outside this range are outliers.

To estimate the extent of adaptation to each environment we competed the evolved populations against an unevolved *lacZ*-marked strain in both heterogeneous and homogeneous environments. Evolved populations from both treatments demonstrated fitness gains relative to the *lacZ* strain (Fig. 3), but there was a significant interaction between evolution and competition environments (likelihood ratio test between models with and without interaction: 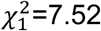, *P* = 0.006; Fig. 3). Evolving in a heterogeneous environment increased relative fitness, heterogeneous-evolved populations competed in heterogeneous environments (relative fitness = 1.90, 95%CI = 1.59-2.22; Table 1) having a significantly higher relative fitness than homogeneous-evolved populations competed in the homogeneous environment (selection rate coefficient = 1.36, 95%CI = 1.04-1.67) (post-hoc contrast between heterogeneous-evolved population in heterogeneous environment *vs*. homogeneous-evolved populations in homogeneous environment: *t-ratio* = −2.56, *d.f.* = 18.7, *P*_*adj*_ = 0.0384). However, this greater adaptation did not transfer into the homogeneous environments: heterogeneous-evolved populations competed in homogeneous environments had lower relative fitness (selection rate coefficient = 1.17, 95%CI = 0.85-1.48) than the same populations competed in the heterogeneous environment (post-hoc contrast between heterogeneous-evolved population in homogeneous *vs*. heterogeneous competition environment: t-ratio = −4.02, d.f. = 10, *P*_*adj*_ = 0.0073). This difference was not observed in the populations evolved in heterogeneous conditions, with no difference in fitness between competition environments (Table 1).

**Table 1.** Results of pairwise comparisons between relative fitness of different populations that differ in evolution and competition environments (either homogeneous or heterogeneous). For each treatment, the evolution environment is followed by the competition environment. *P*-value adjustment: Holm-Bonferroni method for 3 tests. Degrees-of-freedom method: Kenward-Roger.

| contrast | estimate | SE | d.f. | t-ratio | p value |
| --- | --- | --- | --- | --- | --- |
| evolution,competition |  |  |  |  |  |
| homogeneous,homogeneous - homogeneous,heterogeneous | 0.74 | 0.18 | 10.0 | 4.02 | 0.0073 |
| homogeneous,homogeneous - heterogeneous,heterogeneous | 0.55 | 0.21 | 18.7 | 2.56 | 0.0384 |
| heterogeneous,homogeneous - heterogeneous,heterogeneous | -0.03 | 0.18 | 10.0 | -0.15 | 0.8860 |

P value adjustment: Holm-Bonferroni method for 3 tests.

**Figure 3.**
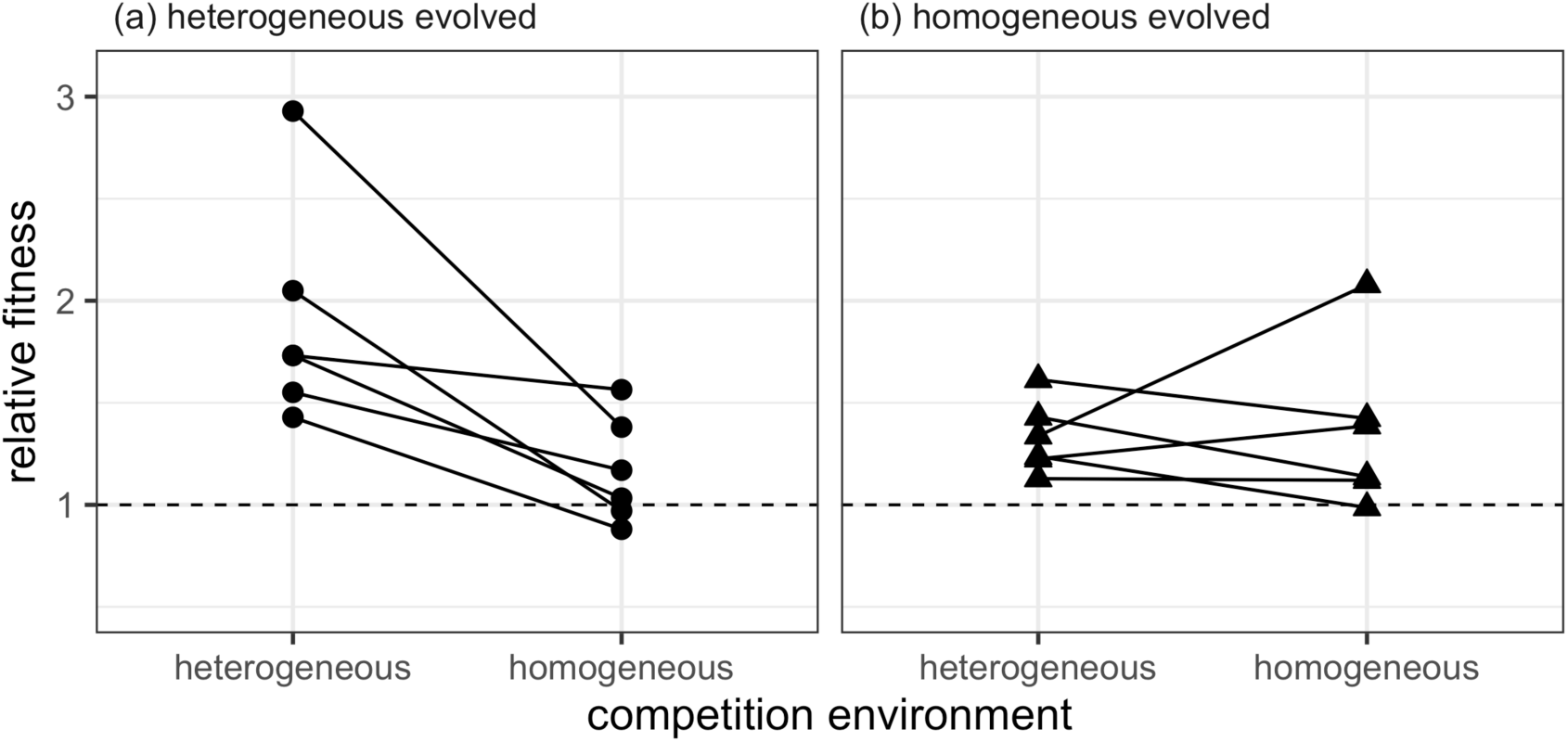
Relative fitness of populations evolved under heterogeneous or homogeneous conditions. Points represent the relative fitness of each population. Lines show the links between each evolved population in each of its competition environments.

### Genomic data

Alongside differences in fitness and phenotypic diversity, we observed some genomic differences between populations evolved in homogeneous and heterogeneous environments (Fig. 4). In terms of genetic distance from the ancestor, heterogeneous populations had a median distance of 0.65 (IQR: 0.53 – 0.7), whereas homogeneous populations had a median distance of 0.35 (IQR: 0.15 – 0.4), but this difference was not significant (Wilcoxon test: W = 6.5, *P* = 0.074; Fig. 4a). However, there were more SNPs / indels in the heterogeneous populations (median = 2.5, IQR = 2-3) compared to those evolved in homogeneous conditions (median = 1, IQR = 1- (Wilcoxon-test: *W* = 4.5, *P* = 0.029; Fig. 4b). Together, this indicates that there was an increased rate of molecular evolution in the heterogeneous populations. Within-population diversity was 0.82 (IQR = 0.81 – 0.85) in heterogeneous populations and 0.45 (IQR = 0.24 – 0.48) in homogeneous populations (Wilcoxon test: W = 5.5, *P* = 0.052; Fig. 4c). Evolution environment (homogeneous *vs*. heterogeneous) significantly altered the genetic composition of the populations (i.e. the Euclidean distance between populations, Fig. 4e, PERMANOVA: *F*_*1,10*_ = 8.92, *R*^*2*^ = 0.47, *P* = 0.0017). This difference was driven in large part by two genetic changes: a SNP in PFLU5698 was observed in all homogeneous populations but never in the heterogeneous populations, and an indel in PFLU1666 was observed in 4 of the 6 heterogeneous populations but never in the homogeneous populations (Fig. 4d). There was no difference in beta-diversity (calculated from distance-to-centroids between groups; Fig. 4e) between homogeneous and heterogeneous populations (homogeneity of multivariate dispersion ANOVA: *F*_*1,10*_ = 3.75, *P* = 0.081).

**Figure 4.**
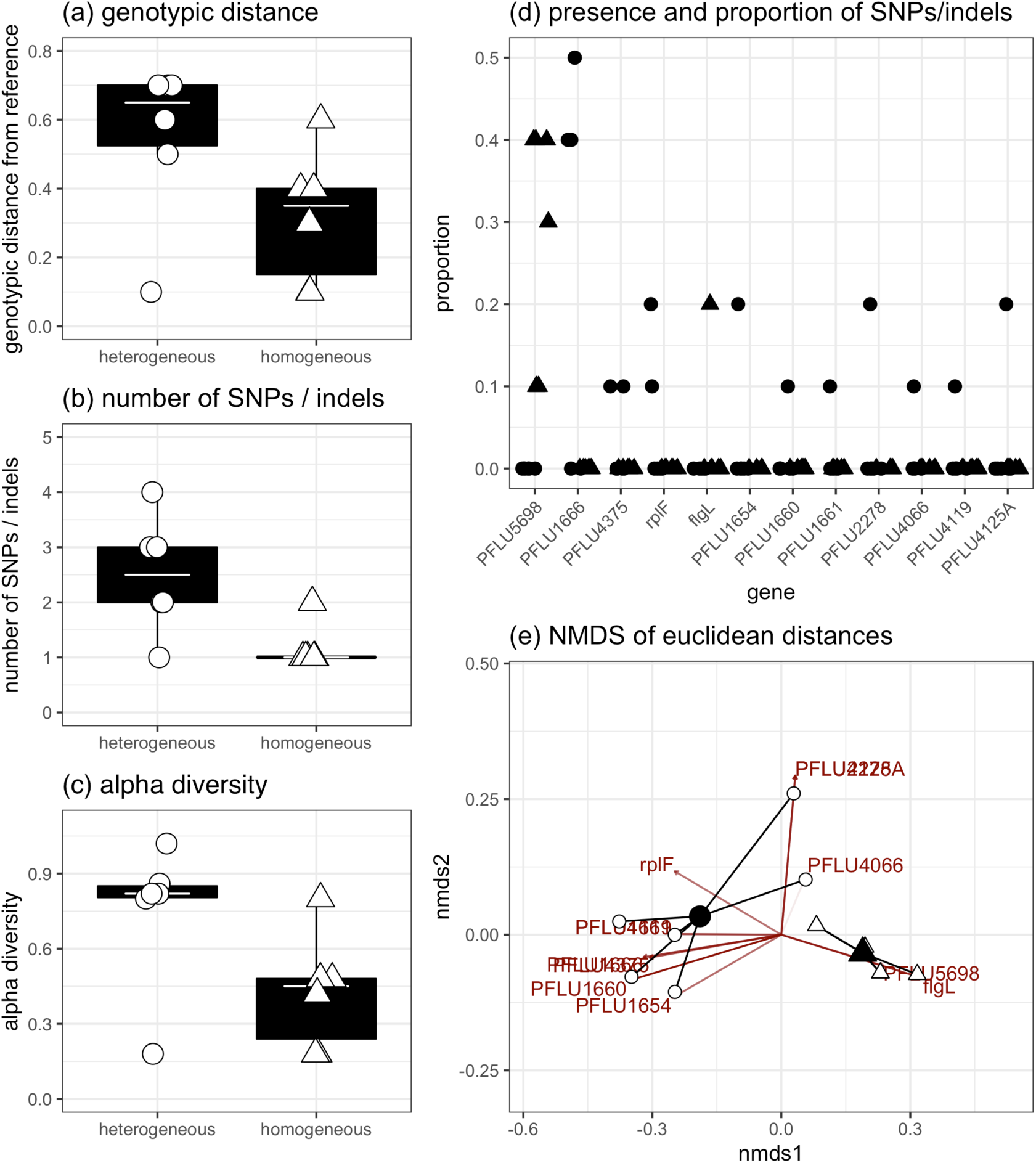
Patterns of genetic differences between populations evolved under heterogeneous or homogeneous conditions. The rate of evolutionary change was estimated using (a) the genetic distance from the ancestor and (b) the number of SNPs. (c) Within-population diversity in homogeneous and heterogeneous populations. (d) Distribution of SNPs and indels across all homogeneous and heterogeneous populations. (e) Non-metric multidimensional scaling (NMDS) plot of Euclidean distance between populations, with centroids (black) and populations (white). In all plots, circles represent heterogeneous populations and triangles are homogeneous populations. In (a-c) tops and bottoms of the bars represent the 75th and 25th percentiles of the data, the white lines are the medians, and the whiskers extend from their respective hinge to the smallest or largest value no further than 1.5 * interquartile range.

## Discussion

Here, we investigated how spatial heterogeneity in an ecologically relevant environment promotes both greater diversification and adaptation of a focal bacterium (*P. fluorescens* SBW25) evolving over 48 days. Consistent with the majority of *in vitro* studies^2-4^ and theoretical work^1,5,7,8^ we show greater phenotypic diversification in heterogeneous potting compost compared with a more homogeneous potting compost-water mix. More importantly, we show that adaptation in the heterogeneous environment is greater than in the homogenous environment – a result that can arise theoretically^16^ and is supported by some, but not all, empirical studies^3,12,17,18^ – suggesting that natural spatial heterogeneity may indeed promote rapid adaptation. The latter result is particularly striking given that population sizes were approximately 3-fold lower in heterogeneous environments, which, if anything, would lead to a reduced mutation supply and less efficient selection, and hence a slower pace of adaptation^30^.

The increased phenotypic diversity in this study is driven, at least partially, by selection, and not simply drift. This is apparent from the greater fitness of the heterogeneously evolved populations in the heterogeneous versus homogeneous environments, which necessarily cannot be explained by drift. Moreover, we have previously shown that *P. fluorescens* genotypes isolated from populations evolved under near-identical conditions that differed in their resource use profiles, as measured using Biolog plates, could reciprocally invade each other from rare^31^. Such negative frequency dependent fitness is a direct indication that diversity is the result of selection^7-9^.

Increased rates of adaptation in spatially heterogeneous environments can theoretically occur because heterogeneity can increase the spatial covariance between genotypes and the local conditions they are best adapted to^15,16^. This relies on the assumption that there are tradeoffs in the ability to use different resources^2^. We are unable to directly test this hypothesis given the complexity of the soil environment and the inability to identify specific resources^31^. An alternative explanation is the greater ability to explore adaptive landscapes because of genetic drift in subdivided populations^13^, but this seems unlikely given the likely role of selection in driving diversification.

The population genomic data are consistent with the phenotypic data. There was evidence for greater rates of molecular evolution, based on the significantly greater numbers of SNPs and indels, in the heterogeneous populations. There was also an indication that within-population diversity was greater for heterogeneous versus homogeneous populations, although the significance of this difference was marginal (*P* = 0.052). While certain genes were mutated across both treatments, different genetic changes were also selected for in the different environments. A SNP in PFLU5698 was observed in all homogeneous populations and resulted in an amino acid change from alanine to valine. While speculative, the protein is 92% similar to di-guanylate cyclase, which has been shown to impact biofilm formation in *Pseudomonas aeruginosa*^32,33^. The other somewhat consistent genetic change was an insertion in PFLU1666, whose predicted function is likely related to fatty acid biosynthesis, which occurred in 4 of the 6 heterogeneous populations. This indel causes a frameshift leading to a truncated protein.

Here, we have shown that phenotypic (and to an extent, genomic) diversification is increased by spatial heterogeneity of an ecologically relevant environment, demonstrating that theoretical predictions and *in vitro* results can be extrapolated to real-world ecological contexts. Moreover, rates of phenotypic and molecular evolution were higher in heterogeneous environments, suggesting that preventing spatial homogenization of environments and populations is key to the emergence and maintenance of diversity, and can potentially promote evolutionary rescue allowing adaptation to anthropogenic environmental change^34^.

## Acknowledgements

This work was funded by NERC. We are grateful to staff at the Centre for Genomic Research, University of Liverpool for technical assistance.

